# Assessment of cladistic data availability for living mammals

**DOI:** 10.1101/022970

**Authors:** Thomas Guillerme, Natalie Cooper

## Abstract

Analyses of living and fossil taxa are crucial for understanding changes in biodiversity through time. The Total Evidence method allows living and fossil taxa to be combined in phylogenies, by using molecular data for living taxa and morphological data for both living and fossil taxa. With this method, substantial overlap of morphological data among living and fossil taxa is crucial for accurately inferring topology. However, although molecular data for living species is widely available, scientists using and generating morphological data mainly focus on fossils. Therefore, there is a gap in our knowledge of neontological morphological data even in well-studied groups such as mammals.

We investigated the amount of morphological (cladistic) data available for living mammals and how this data was phylogenetically distributed across orders. 22 of 28 mammalian orders have *<*25% species with available morphological data; this has implications for the accurate placement of fossil taxa, although the issue is less pronounced at higher taxonomic levels. In most orders, species with available data are randomly distributed across the phylogeny, which may reduce the impact of the problem. We suggest that increased morphological data collection efforts for living taxa are needed to produce accurate Total Evidence phylogenies.

## Introduction

There is an increasing consensus among biologists that studying both living and fossil taxa is essential for fully understanding macroevolutionary patterns and processes [1, 2]. To perform such analyses it is necessary to combine living and fossil taxa in phylogenetic trees. One increasingly popular method, the Total Evidence method [3, 4], combines molecular data from living taxa and morphological data from both living and fossil taxa in a supermatrix (e.g. [5, 4, 6, 1, 7]), producing a phylogeny with living and fossil taxa at the tips. A downside of this method is that it requires molecular data for living taxa and morphological data for both living and fossil taxa. Chunks of this data can be difficult, or impossible, to collect for every taxon in the analysis. For example, fossils rarely have molecular data and incomplete fossil preservation may restrict the amount of morphological data available. Additionally, it has become less common to collect morphological characters for living taxa when molecular data is available (e.g. in [8], only 13% of living taxa have coded morphological data). Unfortunately this missing data can lead to errors in phylogenetic inference. Simulations show that the ability of the Total Evidence method to recover the correct topology decreases when there is little overlap between morphological data in living and fossil taxa, and that the effect of missing data on topology is greatest when living taxa have few morphological data [9]. This is because (1) fossils cannot branch in the correct clade if it contains no morphological data for living taxa; and (2) fossils have a higher probability of branching within clades with more morphological data for living taxa, regardless of whether this is the correct clade [9].

The issues above highlight that it is crucial to have sufficient morphological data for living taxa in a clade before using a Total Evidence approach. However, it is unclear how much morphological data for living taxa is actually available, i.e. already coded from museum specimens and deposited in phylogenetic matrices accessible online, and how this data is distributed across clades. Intuitively, most people assume this kind of data has already been collected, but empirical data suggest otherwise (e.g. in [4, 8, 7]). To investigate this further, we assess the amount of available morphological data for living mammals to determine whether sufficient data exists to build reliable Total Evidence phylogenies in this group. We also determine whether the available cladistic data is phylogenetically overdispersed or clustered across mammalian orders.

## Materials and Mhods

### Data collection and standardisation

We downloaded all cladistic matrices containing any living and/or fossil mammal taxa from three major public databases: MorphoBank (http://www.morphobank.org/ [10]), Graeme Lloyd’s website (graemetlloyd.com/matrmamm.html) and Ross Mounce’s GitHub repository (<u>https://github.com/rossmounce/cladistic-data</u>). We also performed a systematic Google Scholar search for matrices that were not uploaded to these databases (see Supplementary Materials Section 1 for a detailed description of the search procedure). In total, we downloaded 286 matrices containing 5228 unique operational taxonomic units (OTUs). We used OTUs rather than species since entries in the matrices ranged from species to families, and standardised the taxonomy as described in Supplementary Materials (section 1). We designated as “living” all OTUs that were either present in the phylogeny of [11] or the taxonomy of [12].

Matrices with few characters are problematic when comparing available data among matrices because (1) they have less chance of having characters that overlap with those of other matrices [13] and (2) they are more likely to contain a higher proportion of specific characters that are not-applicable across large clades (e.g. “antler ramifications” is a character that is only applicable to Cervidae not all mammals [14]). Therefore we selected only matrices containing *>*100 characters for each OTU. This threshold was chosen to correspond with the number of characters used in [9] and [15]. Results of analyses with no threshold are available in Supplementary Material. After removing matrices with *<*100 characters, we retained 1074 unique living mammal OTUs from 126 matrices.

### Data availability and distribution

To assess the availability of cladistic data for each mammalian order, we calculated the percentage of OTUs with cladistic data at three different taxonomic levels: family, genus and species. We consider orders with *<*25% of living taxa with cladistic data as having low data coverage, and orders with *>*75% of living taxa with cladistic data as having high data coverage.

We investigated whether the available cladistic data for each order was (i) randomly distributed, (ii) overdispersed or (iii) clustered, with respect to phylogeny, using two metrics from community phylogenetics: the Nearest Taxon Index (NTI; [16]) and the Net Relatedness Index (NRI; [16]). NTI is most sensitive to clustering or overdispersion near the tips, whereas NRI is more sensitive to clustering or overdispersion across the whole phylogeny [17]. Both metrics were calculated using the picante package in R [18, 19].

NTI [16] is based on mean nearest neighbour distance (MNND) and is calculated as follows:

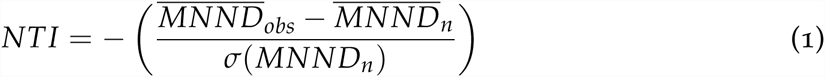

where 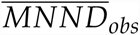 is the observed mean distance between each of *n* taxa with cladistic data and its nearest neighbour with cladistic data in the phylogeny, 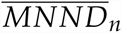 is the mean of 1000 mean MNND between *n* randomly drawn taxa, and *σ*(*MNND*_*n*_) is the standard deviation of these 1000 random MNND values. NRI is calculated in the same way, but MNND is replaced by mean phylogenetic distance (MPD) as follows:

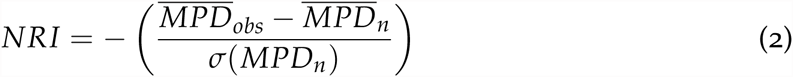
where 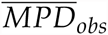 is the observed mean phylogenetic distance of the tree containing only the *n* taxa with cladistic data. Negative NTI and NRI values show that the focal taxa are more overdispersed across the phylogeny than expected by chance, and positive values reflect clustering.

We calculated NTI and NRI values for each mammalian order separately, at each different taxonomic level. For each analysis our focal taxa were those with available cladistic data at that taxonomic level and the phylogeny was that of the order pruned from [11].

## Results

22 of 28 orders have low coverage (*<*25% species with cladistic data) and six have high coverage (*>*75% species with cladistic data) at the species-level. At the genus-level, three orders have low coverage and 12 have high coverage, and at the family-level, no orders have low coverage and 23 have high coverage (Table1).

**Table 1:**
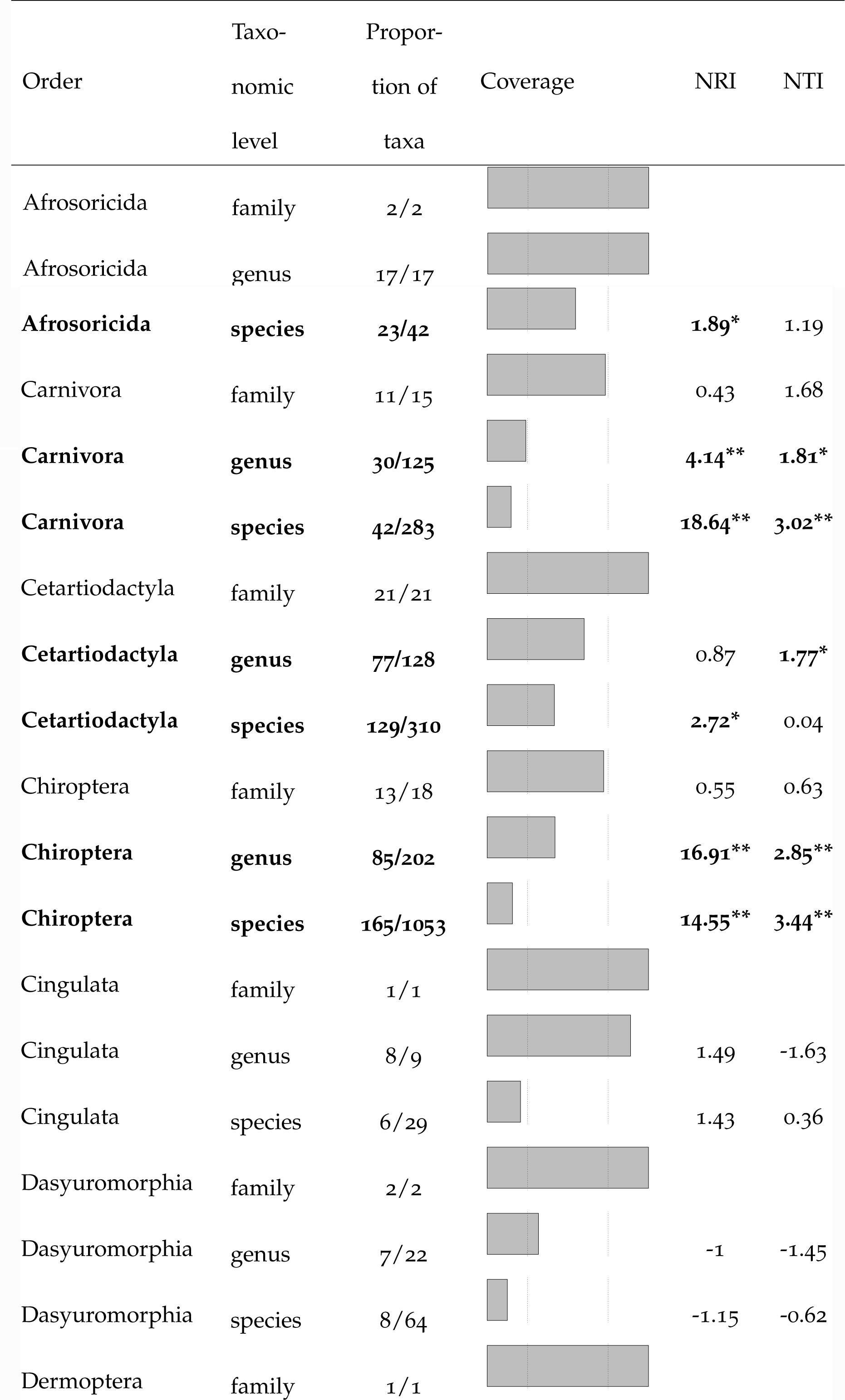
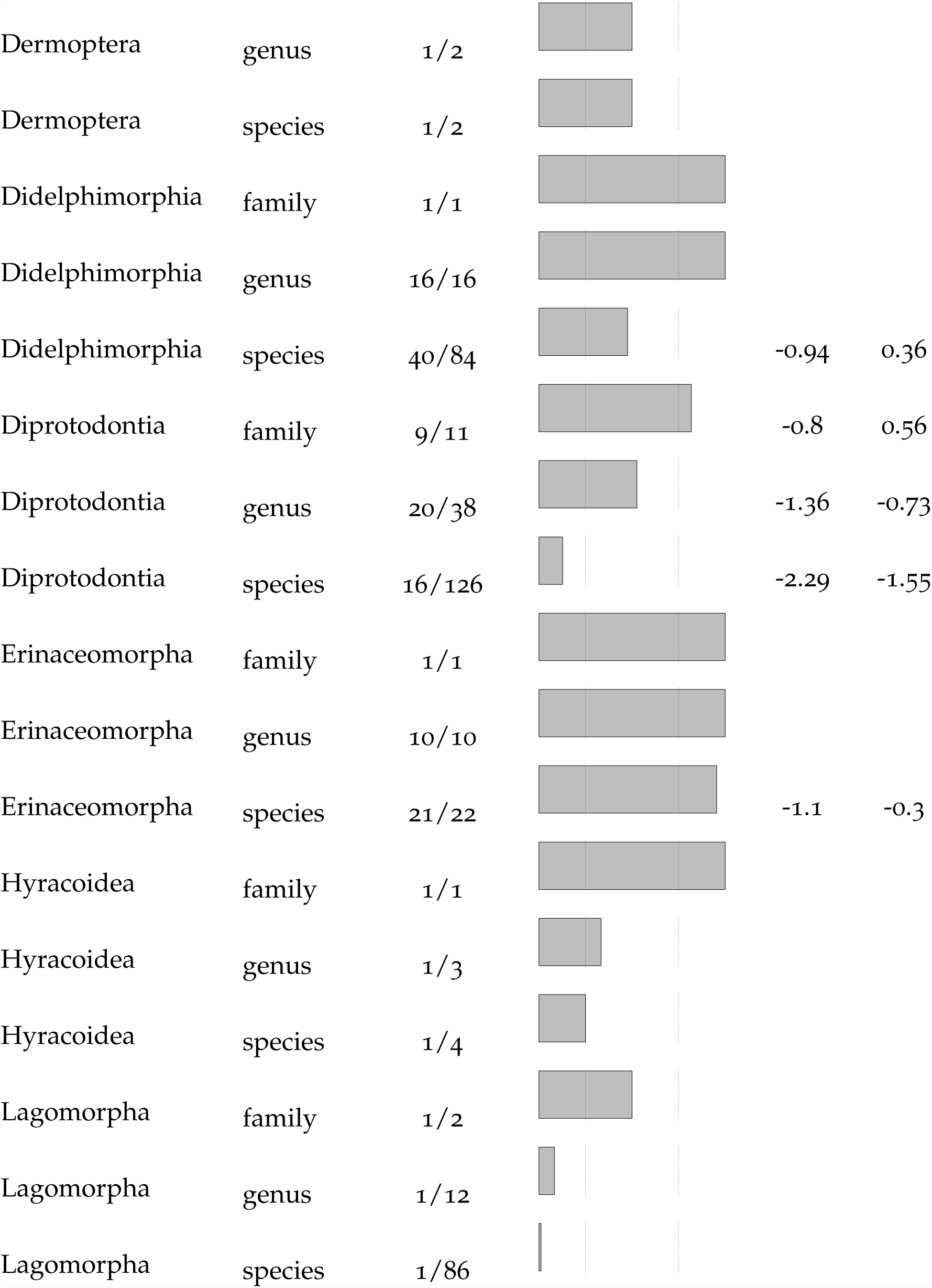
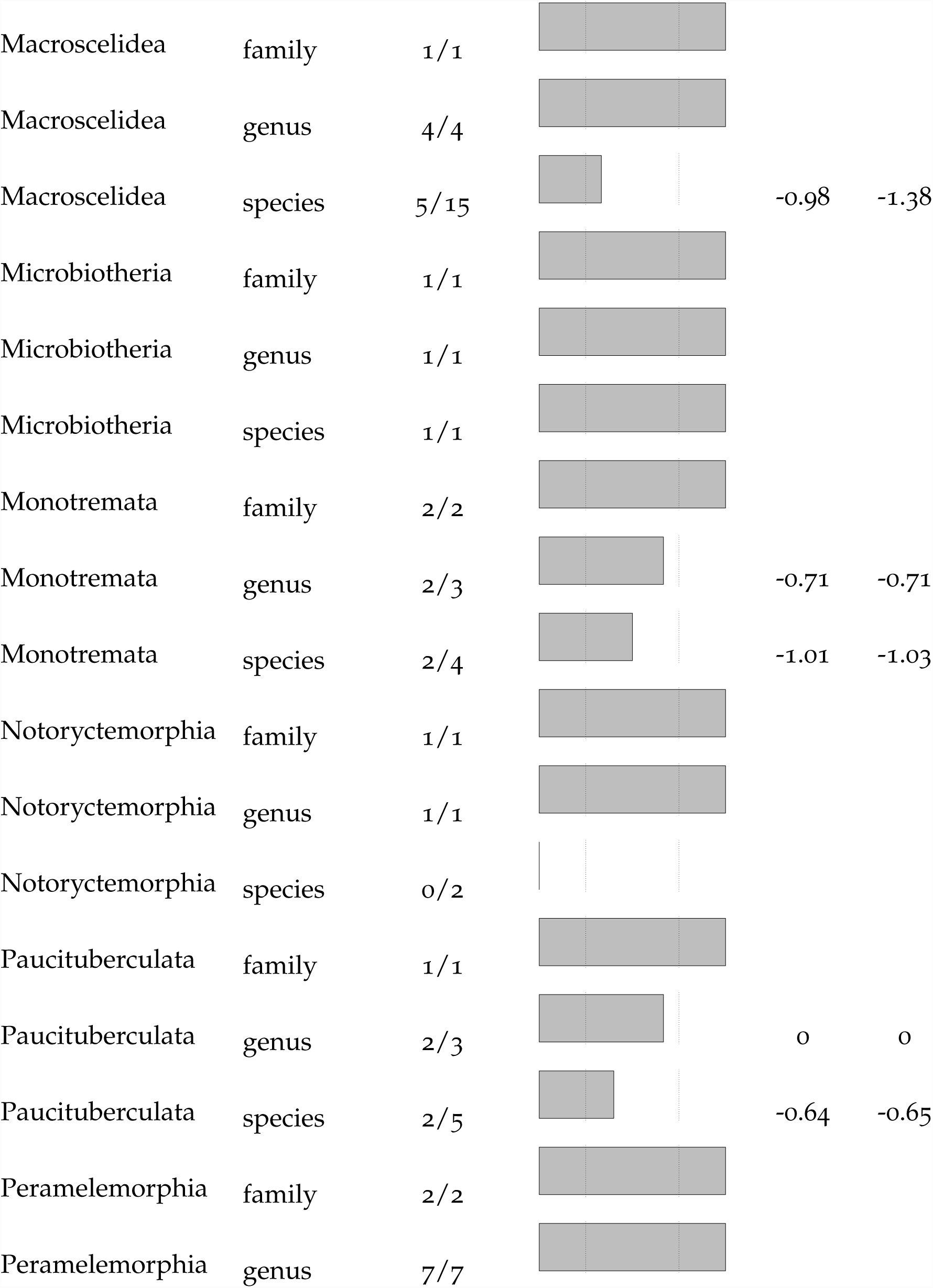
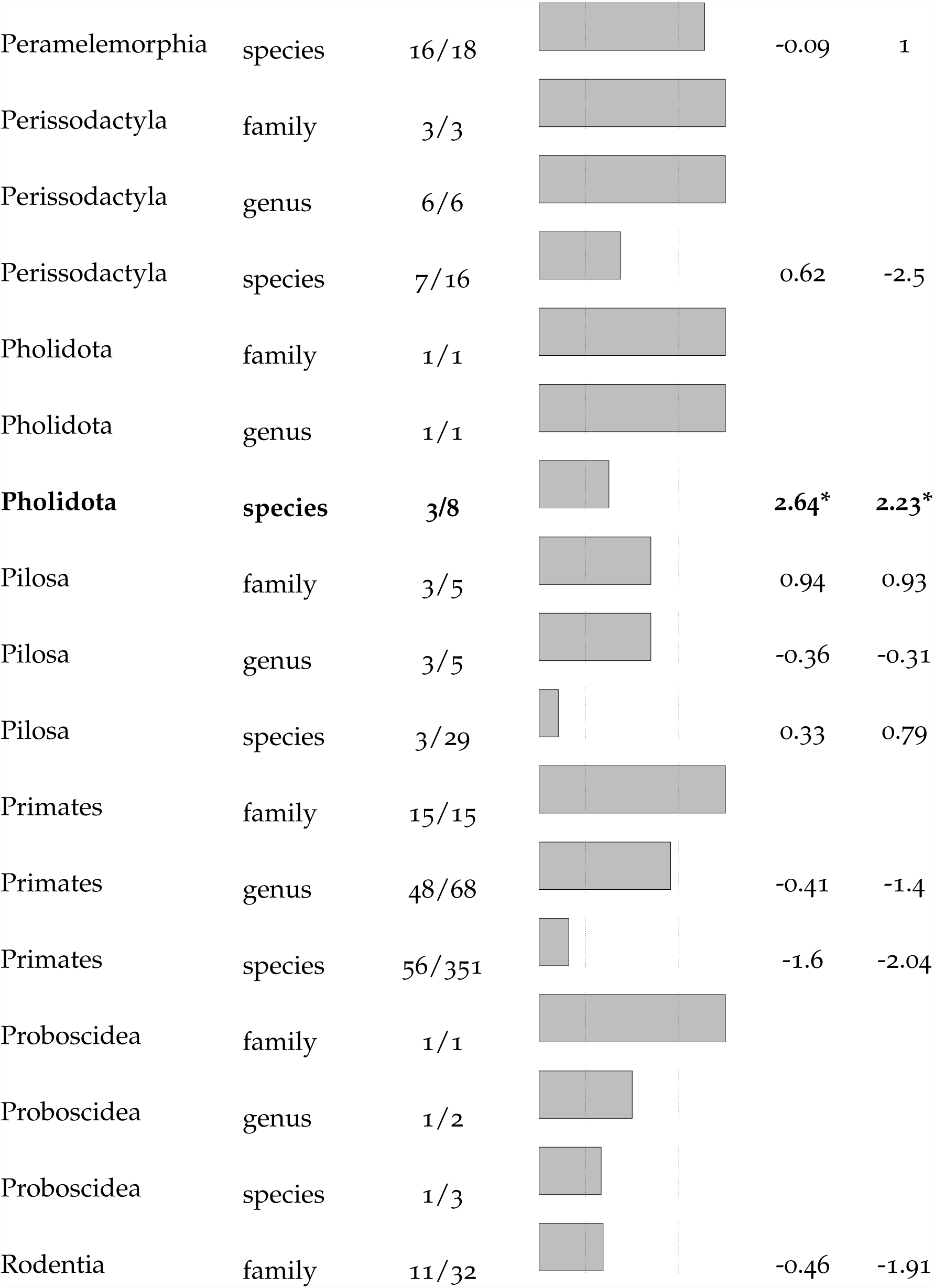
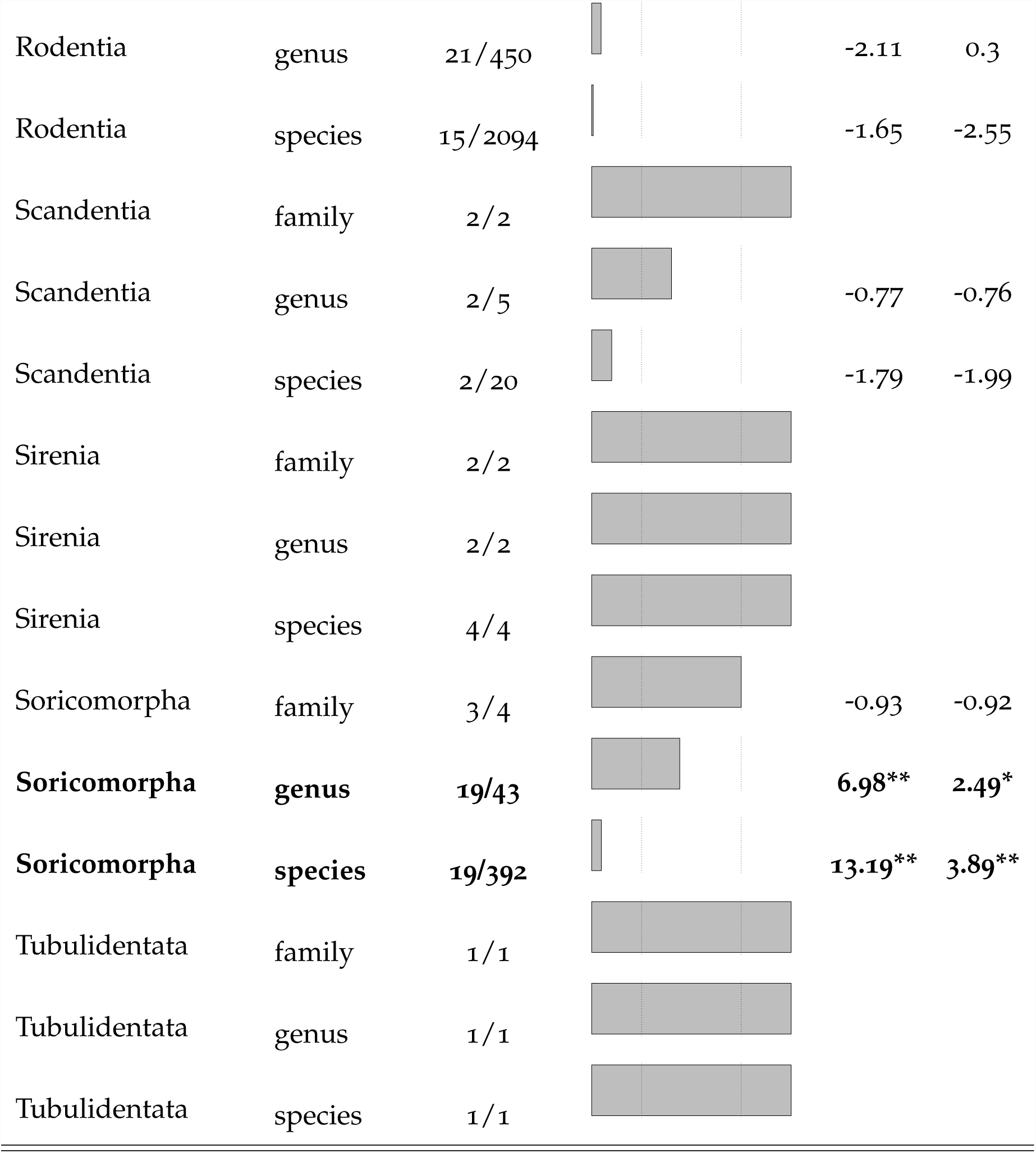
Number of taxa with available cladistic data for mammalian orders at three taxonomic levels. The left vertical bar represents low coverage (*<*25%); the right vertical bar represents high coverage (*>*75%). Negative Net Relatedness Index (NRI) and Nearest Taxon Index (NTI) values indicate phylogenetic overdispersion; positive values indicate phylogenetic clustering. Significant NRI or NTI values are in bold. *p *<*0.05; **p *<*0.01; ***p *<*0.001.

Only six orders had significantly clustered data (Afrosoricida and Pholidota at the species-level, and Carnivora, Cetartiodactyla, Chiroptera and Soricomorpha at both species- and genus-level) and none had significantly overdispersed data (Table 1).

Figure 1 shows randomly distributed OTUs with cladistic data in Primates (Figure 1A) and phylogenetically clustered OTUs with cladistic data in Carnivora (mainly Canidae; Figure 1B).

**Figure 1:**
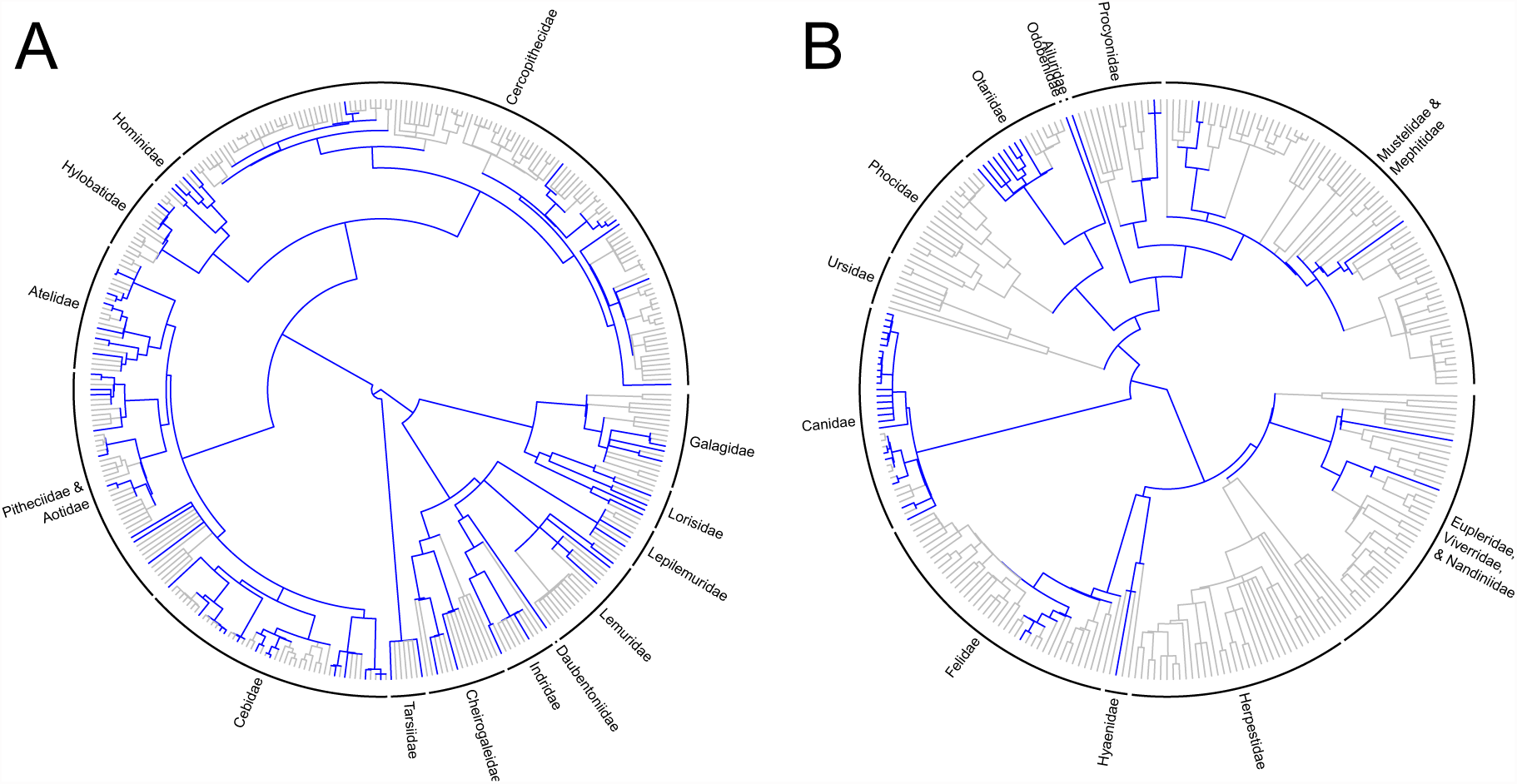
Phylogenetic distribution of species with available cladistic data across two orders (A: Primates; B: Carnivora). Blue branches indicate available cladistic data for the species.

## Discussion

Our results show that although phylogenetic relationships among living mammals are well-resolved (e.g. [11, 20]), most of the data used to build these phylogenies is molecular, and very little cladistic data is available for living mammals compared to fossil mammals (e.g. [21, 22]). This has implications for building Total Evidence phylogenies containing both living and fossil mammals, as without sufficient cladistic data for living species, fossil placements in these trees are very uncertain [9].

The number of living mammalian taxa with no available cladistic data was surprisingly high at the species-level: only six out of 28 orders have a high coverage of taxa with available cladistic data. This high coverage threshold of 75% of taxa with available cladistic data represents the minimum amount of data required before missing data has a significant effect on the topology of Total Evidence trees [9]. Beyond this threshold, there is considerable displacement of wildcard taxa (*sensu* [23]) and decreased clade conservation [9]. Therefore we expect difficulties in placement of fossil taxa at the species-level in most mammalian orders, but fewer issues at higher taxonomic levels. This point is important from a practical point of view because of the slight discrepancy between neontological and palaeontological species concepts. While neontological species are described using morphology, genes, distribution etc.; palaeontological species can be based only on morphological, spatial and temporal data (e.g. [22]). Therefore, most palaeontological studies use genus as their smallest OTU (e.g. [22, 21]), so data availability at the genus-level in living mammals should be our primary concern when building phylogenies of living and fossil taxa.

When few species have available cladistic data, the ideal scenario is for them to be phylogenetically overdispersed to maximize the possibilities of a fossil branching from the right clade. The second best scenario is that species with cladistic data are randomly distributed across the phylogeny. Here we expect no special bias in the placement of fossils [9], it is therefore encouraging that for most orders, species with cladistic data were randomly distributed across the phylogeny. The worst case scenario for fossil placement is that species with cladistic data are phylogenetically clustered. Then we expect two major biases to occur: first, fossils will not be able to branch within a clade containing no data, and second, fossils will have higher probability of branching within the most sampled clade by chance. Our results suggest that this may be problematic at the genus-level in Carnivora, Cetartiodactyla, Chiroptera and Soricomorpha. For example, a Carnivora fossil will be unable to branch in the Herpestidae, and will have more chance to randomly branch within Canidae (Figure 1B).

Despite the absence of good cladistic data coverage for living mammals, the Total Evidence method still seems to be the most promising way of combining living and fossil data for macroevolutionary analyses. Following the recommendations in [9], we need to code cladistic characters for as many living species possible. Fortunately, data for living mammals is usually readily available in natural history collections, therefore, we propose that an increased effort be put into coding morphological characters from living species, possibly by engaging in collaborative data collection projects. Such an effort would be valuable not only to phylogeneticists, but also to any researcher focusing understanding macroevolutionary patterns and processes.

## Ethics statement

N/A

## Data accessibility statement

All data and analysis code is available on GitHub

(<u>https://github.com/TGuillerme/Missing_living_mammals</u>).

## Authors’ Contributions

T.G. and N.C conceived and designed the experiments. T.G. performed the experiments and analysed the data. T.G. and N.C. contributed to the writing of the manuscript. All authors approved the final version of the manuscript.

## Competing Interests

We have no competing interests.

## Acknowledgments

We thank David Bapst, Graeme Lloyd, Nick Matzke and April Wright.

## Funding statement

This work was funded by a European Commission CORDIS Seventh Framework Programme (FP7) Marie Curie CIG grant (proposal number: 321696).

